# Damselflies Overcome Color Saturation Barriers of Photonic Glasses *via* Structural Dispersion and Pigment Loading

**DOI:** 10.64898/2025.11.29.691096

**Authors:** Tali Lemcoff, Lotem Alus, Albert Batushansky, Yahel Fishman, Nila Theodor, Keshet Shavit, Lahav Hyitner, Almut Kelber, Johannes S. Haataja, Dan Oron, Benjamin A. Palmer

## Abstract

Biology’s strategies for manipulating light offer rich inspiration for the design of sustainable replacements to conventional pigments in paints, coatings, and displays. For these applications, where angle-independent color is required, photonic glasses, composed of random arrangements of dielectric spheres, offer a promising solution. However, their intrinsic disorder, particularly from particle polydispersity, fundamentally limits their color saturation and practical utility. In contrast, insects like damselflies and dragonflies exhibit surprisingly vivid, non-iridescent structural colors, despite relying on disordered photonic structures. Here, we show how damselflies combine compositional and structural dispersion to overcome color saturation limits of photonic glasses. Firstly, doping transparent particles with yellow pigments dramatically enhances blue-green structural resonances by the coupled effects of narrowband absorption and refractive index (material) dispersion. Secondly, the refractive index of the nanospheres varies with their size and crystallinity. This gives rise to a ‘structural dispersion’ which maintains consistent optical path lengths in polydisperse assemblies, preserving high color purity. Finally, we show how damselflies tune their structural colors during maturation by precisely modulating the size of the nanospheres. Remarkably, the tuning of particle size, refractive index and pigment loading, arises naturally during the development of the pigment cells – where the pteridine nanospheres undergo a process of densification, crystallization and metabolic maturation.

## Introduction

From iridescent butterflies (1–3) to ultra-white beetles (4) and shrimp (5), biology’s optical systems have revealed new fundamental principles in scattering physics which inspire the development of sustainable optical materials. Bioinspiration is becoming increasingly important as concerns grow over the toxicity and environmental impact of conventional pigments in foods, cosmetics, pharmaceuticals and textiles (6–9). While numerous iridescent biomimetic materials have been produced (10–13), angle-independent colors are required for many applications (14, 15). Photonic glasses, composed of random arrangements of dielectric spheres, offer a viable solution. Here, non-iridescent color emerges from the combination of short-range order, resonant scattering and possibly absorption (16–19). However, the inherent disorder in these systems, particularly the size distribution of scattering spheres, typically results in low color saturation (20–23). Several strategies for enhancing saturation have been proposed, including the use of core-shell particles (23, 24), control of layer thickness and packing density (25, 26), dual-size particle systems (25, 27) and the incorporation of narrowband (28) or broadband (20, 29–33) absorbers.

Although few photonic glasses in animals have been reported (34–39), a striking case is found in blue-tailed damselflies (*Ischnura elegans*) (40–44). During the maturation of juvenile males, the color of the thorax changes from vivid green to blue (42, 45). The highly saturated structural colors are produced by arrays of nanospheres (40–43), thought to be composed of pteridines (42, 44), contained within epidermal chromatophore cells. Underlying this, melanin/ommochrome pigment cells likely enhance contrast by absorbing stray light (42, 44). Prum *et al*. (41) showed that blue structural colors of Odonata (e.g., dragonflies and damselflies) are produced from coherent scattering by the nanospheres, rather than Tyndall scattering and Henze *et al*. (42) proposed that changes in the relative concentrations of xanthopterin and erythropterin pigments may be responsible for tuning the coloration.

Here, we show that damselflies precisely modulate the size of pteridine nanospheres to tune the color of a photonic glass and utilize two surprising strategies to dramatically enhance color saturation. Firstly, doping of transparent pteridine nanospheres with xanthopterin, a yellow pigment, enhances structural resonances by the coupled effects of off-resonance absorbance and refractive index (material) dispersion. Secondly, the refractive index of the nanospheres is found to be linearly dependent on their size and crystallinity. This gives rise to a type of ‘structural dispersion’ which imparts the system with a tolerance against reflectance broadening caused by particle polydispersity – maintaining its high color saturation. The modulation of particle size, refractive index and pigment loading are all consequences of the natural development of the chromatophores – where the pteridine nanospheres undergo a process of formation, densification, crystallization and metabolic maturation during ontogeny.

## Results

### Ontogenetic color change and ultrastructure of the damselfly chromatophore cells

Reflectance spectra from the thoraxes of over 100 male *I. elegans* damselflies (Fig 1A-B), exhibit a reflectance peak shift from 530 to 510 nm as they transform from green to blue during maturation (Fig. 1C i-iv). The blue tail exhibits a reflectance peak at 470 nm (Fig. 1C, Tail) which remains unchanged throughout ontogeny. Conversion of these spectra into CIE color space (46) (Fig. 1D) shows that the damselfly colors are highly saturated (close to the boundary of the most saturated colors in the sRGB color space) and undergo a gradual blue-shift during maturation (Fig. 1E). We note that the CIE color space quantitatively describes colors based on human perception and excludes the ultraviolet range which is relevant for insect and bird vision.

**Figure 1.**
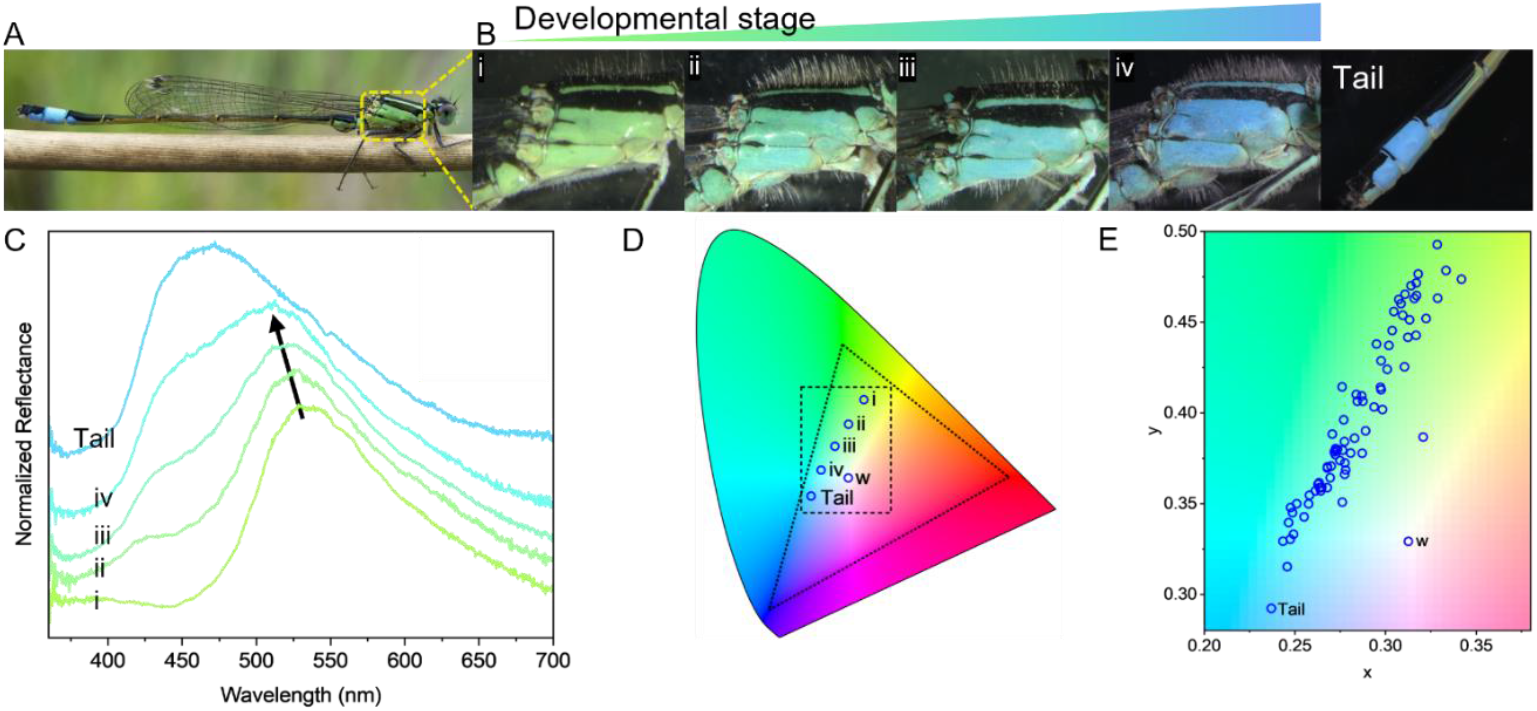
**(A)** The male blue-tailed damselfly (*Ischnura elegans*). Photo credit: Ian Kirk. **(B)** Juvenile thoraxes during the development (i-iv). Right; tail region. **(C)** Reflectance spectra from the thoraxes and tail also exhibiting an intensifying shoulder peak at 430 nm. Black arrow indicates the color progression over ontogeny. **(D)** The spectra in (C) plotted on a CIE chromaticity diagram. w; white point of standard illuminant D65 corresponding to average daylight (saturation increases with distance from the white point). Dotted triangle; the sRGB color space, dashed rectangle; enlarged region of interest in (E). **(E)** Reflectance spectra of ∼100 damselflies plotted on a CIE chromaticity diagram, exhibiting a gradual color change from green to blue.

Cryogenic scanning electron microscopy (cryo-SEM) reveals that the ultrastructure of the thorax remains similar throughout development – with a random or quasi-ordered array of membrane-bound nanospheres (resembling a photonic glass (16, 17, 47)) in the distal epidermis, overlying larger pigment granules in the proximal epidermis (Fig. 2A and SI Appendix, Fig. S1). However, as the damselflies transition from green to blue, the average size of nanospheres in the thorax decreases from 339 to 306 nm, whilst the nanosphere size in the tails remains constant at ∼282 nm (SD = 4-8.6% across all samples) (Fig. 2B, C). The filling fraction of nanospheres was estimated to be 46 ± 5% in the thorax and 54 ± 5% in the tail.

**Figure 2.**
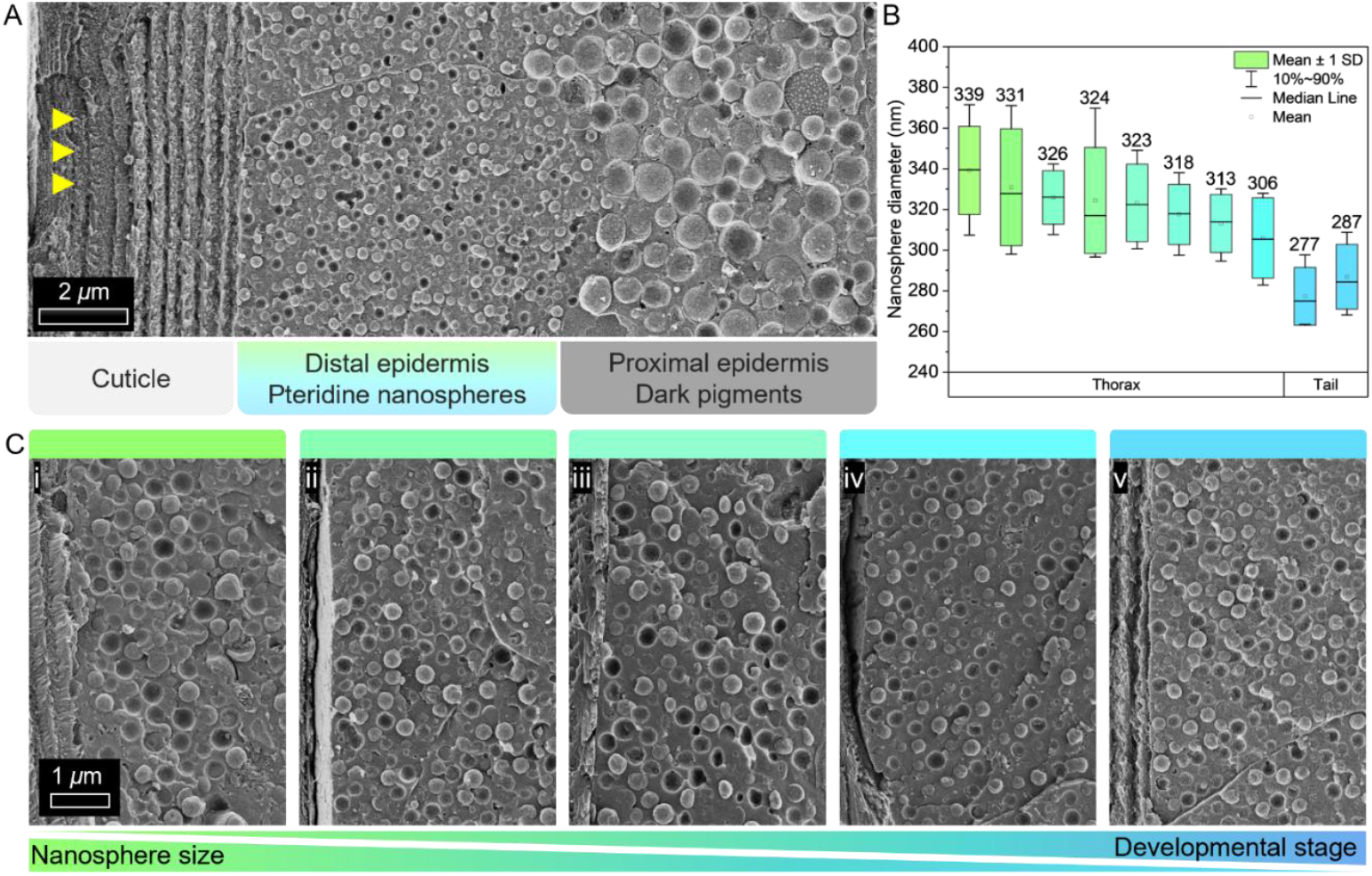
**(A)** Cryo-SEM micrograph of a damselfly tail showing a cross-section through the epidermis. Yellow arrowheads; direction of incident light. **(B)** Box chart showing the sizes of nanospheres extracted from individuals with different thorax colors as well as tails. The average diameter measured in each specimen (nm) is displayed on the chart. **(C)** Cryo-SEM images of pteridine nanospheres in the distal epidermis in thoraxes (i-iv) and a tail (v). The colors of the box chart in (B) and panel bars in (C) are the CIE representation of the reflectance measured from the specific tail/thorax analyzed.

### Structural and optical properties of nanospheres

At the early stages of maturation, when the thorax is green, the nanospheres display a smooth or slightly granulated texture (Fig. 3Ai and SI Appendix, Fig. S2; extended sample size) which becomes more pronounced as development progresses (Fig. 3Aii). Later in development, the granules arrange into radial spokes (white arrowheads, Fig. 3), initially occupying the center of the nanosphere, with the edge containing an amorphous material (white brackets, Fig. 3Aii-iii). Densification of the spokes occurs concomitantly with a contraction of the nanosphere membrane (white arrows, Fig. 3A) and a decrease in nanosphere diameter (Fig. 3Aii-iv). Finally, the nanospheres undergo a further decrease in size as they transform from a spoke-like to onion-like texture (Fig. 3Av) - resembling crystalline isoxanthopterin nanospheres in crustacean visual systems (34, 48, 49). These morphological changes indicate that the nanospheres undergo a process of formation during damselfly maturation – likely related to an ongoing crystallization process. Electron diffraction measurements confirm this. In early juveniles, the nanospheres exhibit an extremely weak, single diffraction ring with a d spacing of ∼3.2 Å – associated with stacking of pteridine molecules (Fig. 3B) (5, 50). As the damselflies mature, the diffraction intensity of the nanospheres increases (Fig. 3B and SI Appendix, Fig. S3). In agreement with textural observations (Fig. 3A), this indicates that the crystallinity and density of the nanospheres increase during maturation, causing a reduction in their volume. Selected area electron diffraction acquired from different locations along the nanosphere edge exhibit diffraction arcs (Fig. 3C), whose azimuthal angle demonstrates the nanospheres are composed of 1-dimensionally ordered, stacked molecular assemblies projecting radially outwards from the sphere. This creates a spherulitic nanosphere (commensurate with the ‘spoke’ like texture observed in the cryo-SEM images, Fig. 3A) closely resembling isoxanthopterin spherulites in cleaner shrimp (5) and uric acid spherulites in fish (50).

**Figure 3.**
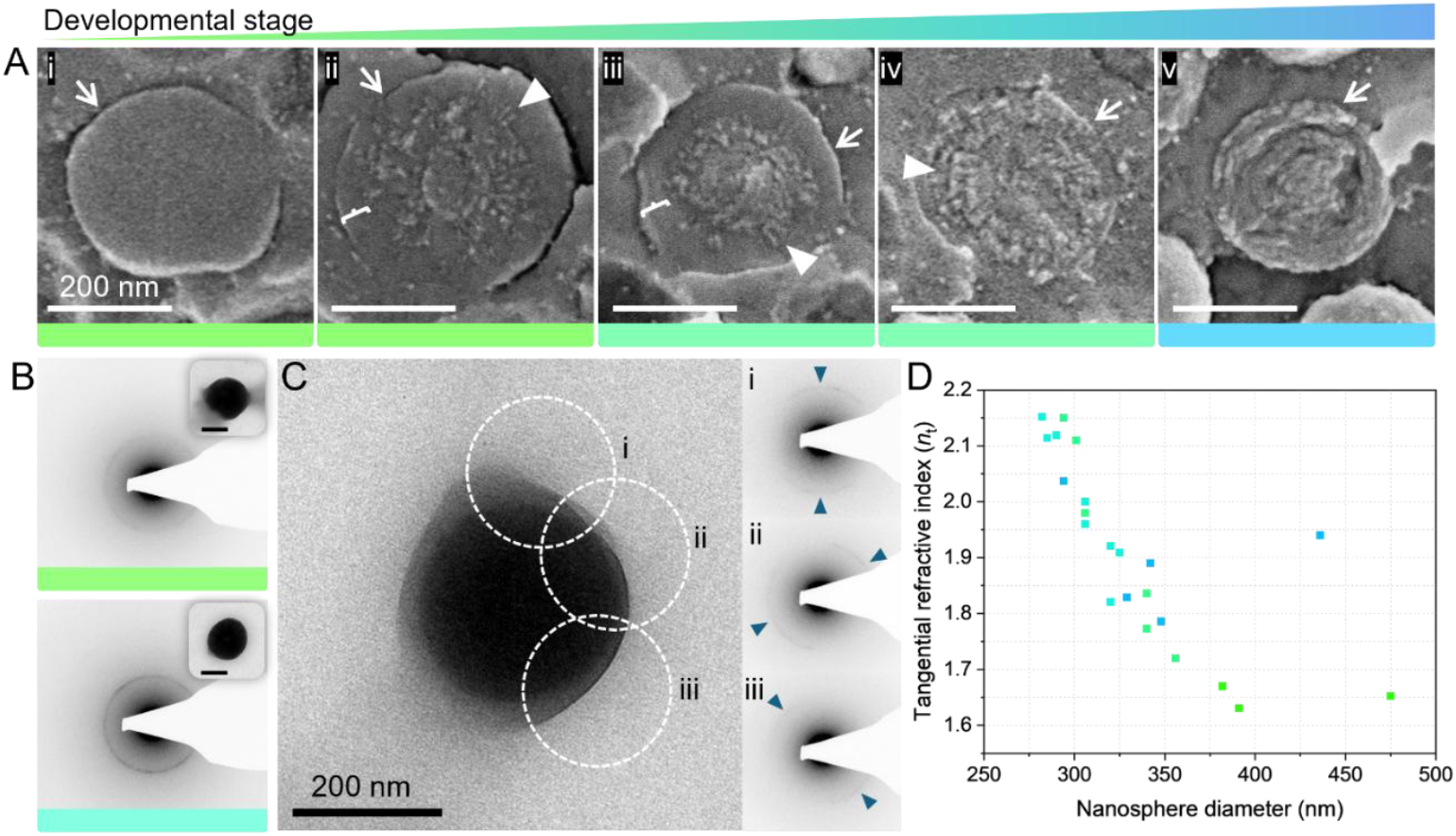
**(A)** Cryo-SEM micrographs nanosphere cross-sections exhibiting different stages of formation during ontogeny. Arrowheads; radial spokes, brackets; layer of amorphous material, arrows; nanosphere limiting membrane. **(B)** Electron diffraction from nanospheres extracted from (top) a damselfly early in development and (bottom) later in development. Scale bars; 200 nm. **(C)** TEM micrograph and corresponding selected area electron diffraction performed on different locations along the nanosphere edge exhibiting diffraction arcs (arrowheads). **(D)** Scatter plot showing the measured tangential refractive index (*n*_t_) of nanospheres with varying diameters. Color panels (A, B) and colored data points (D); CIE representation of the reflectance measured from the specific tail/thorax analyzed.

We note that the intensity of the stacking reflection remains relatively weak at all stages of damselfly development compared to that in cleaner shrimp, indicating they are generally less crystalline (5). In previous work (5), the spherulitic arrangement of isoxanthopterin molecules produces extremely birefringent nanospheres with a high tangential refractive index (n = 1.96) but a low radial refractive index (n = 1.4) (48, 49, 51, 52). Birefringence results from the preferred orientation of the planar isoxanthopterin molecules, which have a highly anisotropic molecular polarizability (53). The degree of birefringence is dictated by the degree of molecular alignment and thus, crystallinity. In damselflies, poorly crystalline nanospheres (in early juveniles) are expected to be weakly birefringent (with a tangential refractive index close to the weighted average of the refractive indices of the component pteridines). As development progresses and crystallinity increases, both the birefringence and the tangential refractive index of the nanospheres are also expected to increase.

To test this premise, we measured the refractive index of individual nanospheres extracted from differently colored damselflies using a scattering method described by Beck *et al*. (51). The measurements revealed an intriguing near-linear correlation between the tangential refractive index (*n*t) of individual nanospheres and their diameters. As the nanosphere size decreases from 391 nm to 282 nm, their tangential refractive index at the resonant scattering wavelength increases from 1.63 (corresponding to a isotropic sphere) to well above 2 (corresponding to a nearly fully crystalline one) (Fig. 3D). This behavior is dominated by the increasing ordering and alignment of pteridine molecules in the nanospheres as they crystallize during chromatophore development and has some contribution from material dispersion (as the resonance blue shits for smaller spheres).

### Correlation between pteridine pigment abundance and thorax color

To investigate whether there is a correlation between thorax color and the pteridine content of the nanospheres, we quantified pteridine concentrations from the distal epidermis of 45 damselflies using liquid chromatography-mass spectrometry (LC-MS). K-means clustering was used to group the samples based on the concentrations of the 7 most abundant pteridines, yielding two clusters, representing groups of individuals with a similar pattern of pteridine abundance (Fig. 4A). The reflectance spectra from the same thoraxes are plotted on a CIE diagram, with the color of the data points (red or blue) denoting their cluster classification (Fig. 4B). Cluster 1 predominantly incorporates green samples, and cluster 2, the rest of the samples. The most abundant pteridines in both clusters are leucopterin (colorless), xanthopterin (yellow pigment) and isoxanthopterin (colorless) (Fig. 4C). The relative amount of yellow pigment (xanthopterin), compared to the colorless pteridines, decreases from 29% in green thoraxes to 23% in blue thoraxes, concomitant with a relative increase in leucopterin from 48 to 61% (Fig. 4D). The same chemical analysis was performed on tails from the same individuals (Fig. 4E), which remain blue throughout their lifetime and thus serve as an internal control. Similarly to the thoraxes, as damselflies mature, the relative concentration of xanthopterin decreases from 30 to 24% as leucopterin increases from 58 to 67% (Fig. 4D). This indicates that changes in the relative abundance of yellow-to-colorless pteridines during maturation do not account for the ontogenetic color change. However, the relative increase in leucopterin and decrease in xanthopterin and 7,8-dihydroxanthopterin (Fig. 4E) in the thoraxes and tails suggests a maturing metabolic process where precursors, xanthopterin and 7,8-dihydroxanthopterin are converted enzymatically (*via* xanthine dehydrogenase) into the less soluble leucopterin – an end product of the pteridine biosynthetic pathway in insects (54–56).

**Figure 4.**
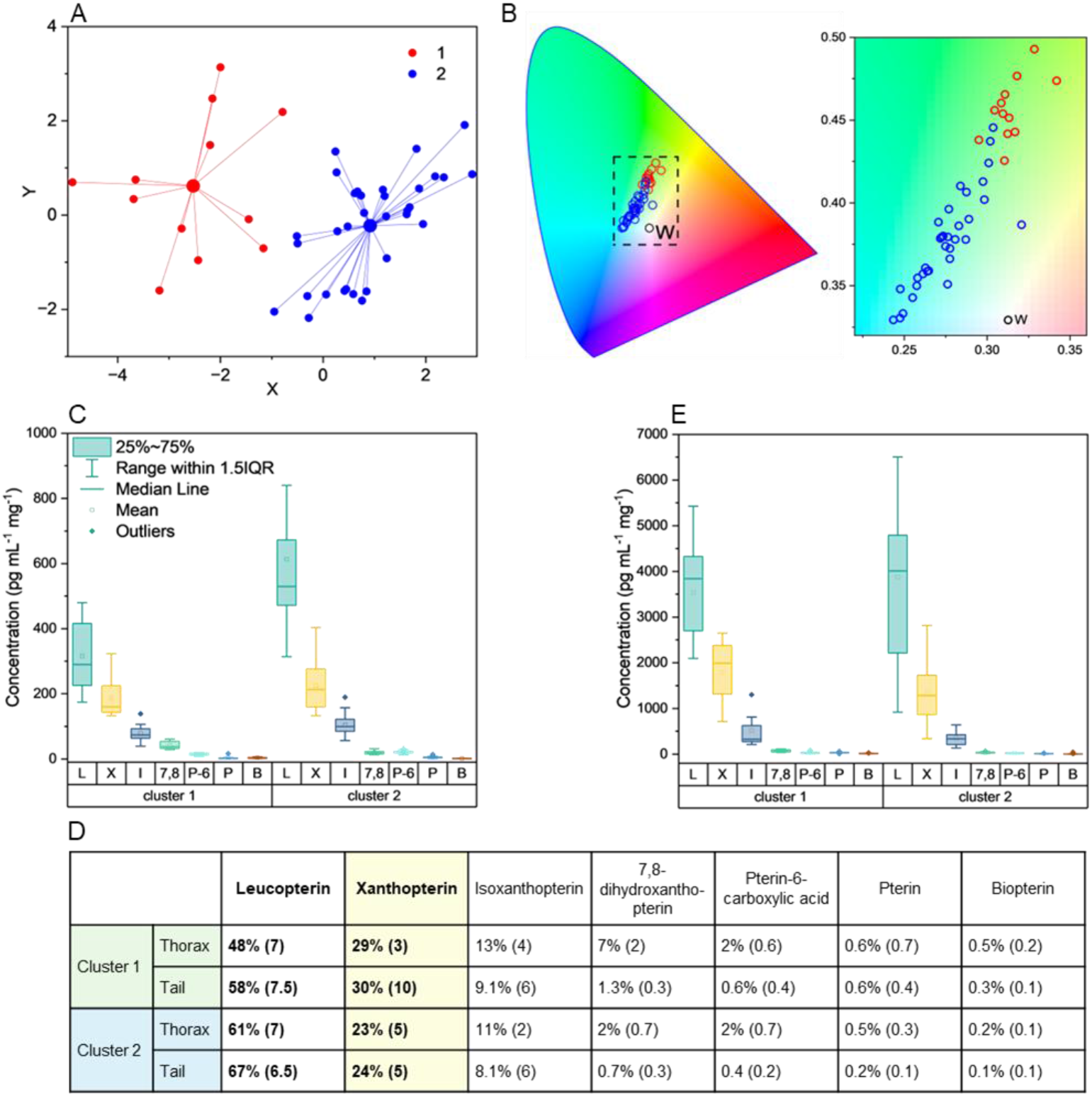
**(A)** K-means clustering of pteridine concentration data from damselflies of different colors. Each cluster represents a group of individuals with a similar pattern of pteridine concentrations. **(B)** CIE chromaticity diagram denoting reflectance spectra from the thoraxes of each damselfly in the analysis. Red and blue data points; classification to clusters 1 and 2 from (A). w; white point of standard illuminant D65 corresponding to average daylight. **(C)** Average concentration (pg mL^-1^ mg^-1^ of tissue) of pteridines in damselfly thoraxes of each of the clusters in (A). (7,8; 7,8-dihydroxanthopterin, B; biopterin, I; isoxanthopterin, L; leucopterin; P; pterin, P-6; pterin-6-carboxylic acid, X; xanthopterin). **(D)** Relative abundance of pteridines in thoraxes and tails of each cluster (percent standard deviation). **(E)** Average concentrations (pg mL^-1^ mg^-1^ of tissue) of pteridines in damselfly tails of each of the clusters in (A).

Important observations are made when considering the absolute abundance (per mass of tissue) of pteridines in the samples (Fig. 4C). When the thoraxes transition from green (cluster 1) to blue (cluster 2), the absolute abundance of leucopterin, xanthopterin and isoxanthopterin increase by 94%, 33%, and 21%, respectively (the mass of tissue analyzed in the specimens of the two groups does not differ significantly). The increase in pteridine abundance in the thorax during maturation indicates the constituent pteridine nanosphere organelles are in a state of high metabolic activity associated with their formation. The increase in absolute pteridine abundance provides an underlying metabolic cause of the densification and crystallization of the nanospheres in the thorax during maturation (Fig. 3). In contrast, in the tails, the absolute abundance of leucopterin increases by only 10%, while xanthopterin and isoxanthopterin decrease by 26% and 33% from cluster 1 to cluster 2. This suggests the nanospheres in the tail may have already attained a more mature and thus dense, crystalline state.

Taking the morphological, structural (Fig. 2, 3) and compositional (Fig. 4) observations together, the ontogenetic color change in damselflies is associated with a reduction in nanosphere size which appears to be caused by densification of pteridines during crystallization. This crystallization process is correlated to an increase in the absolute pteridine content of the nanospheres and in their tangential refractive index (*nt*). This interpretation is consistent with the observation of Hinnekint et al., (45) who reported that damselfly color changes during maturation are correlated with the dehydration of the tissue – another possible driver of densification/crystallization.

### Optical properties of pigment-loaded nanosphere assemblies

Photonic glasses typically produce low color saturation due to multiple scattering effects (20–23). To rationalize how damselflies produce and tune highly saturated colors, we used molecular dynamics (MD) simulations to generate random particle assemblies and the finite difference time domain (FDTD) method to simulate their reflectance spectra. Simulations were performed using experimentally measured nanosphere sizes, with a 5% polydispersity in diameter and a 55% filling fraction. Initially, the nanospheres were modeled as transparent spheres, with an isotropic refractive index - a simple approximation used previously for non-absorbing dielectric materials such as colorless pteridines (5, 34). The refractive indices of the particles were made linearly dependent on size based on our experimental measurements (Fig. 3D and SI Appendix, Fig. S4).

Simulations reveal a structural peak in the visible which blue-shifts from ∼550 to ∼515 nm as the nanosphere size decreases from 340 to 280 nm (Fig. 5A). The peak positions closely match those observed experimentally, albeit with a slight red-shift (∼20 nm) likely resulting from a small overestimation of the nanospheres’ refractive index (elaborated on in the Methods section). The shorter wavelength reflectance observed in the tail compared to thoraxes may be due to the higher nanosphere packing density in the tails. These results show that the green-to-blue ontogenetic color shift is driven by a decrease in nanosphere size. However, while the experimental data show low reflectance from 300–450 nm and high color saturation (s = 0.4–0.7), this simulation yielded a more broadband spectral response, with relatively high reflectance throughout the visible, resulting in low color saturation (s = 0.08–0.14). This is likely because the simulation assumes the particles are transparent, while many pteridines absorb in the UV-A (315-400 nm). We therefore incorporated the wavelength-dependent real (*n*) and imaginary (κ, extinction coefficient) components of the complex refractive index (*n*+iκ) of leucopterin (the most abundant molecule in the nanospheres) into the simulation (Fig. 5B). The extinction coefficient (κ) was calculated from a scaled absorbance measurement (eq. 1, SI Appendix, Fig. S5A-B). The real part of the refractive index (*n*) was computed *via* the Kramers–Kronig relation (57, 58) (eq. 2, SI Appendix, Fig. S5A), with the measured tangential refractive index (*n*_t_) used as the refractive index at infinite wavelength (*n*∞) to account for the refractive index size dependence (SI Appendix, Fig. S4). The resulting simulations reveal similar reflectance peaks in the visible (with a shoulder peak at 460 nm) but with very low reflectance between 300-400 nm due to leucopterin’s UV absorbance and refractive index dispersion (SI Appendix, Fig. S5A, S6A). Consequently, the spectra more closely match the experimental reflectance, and the reflected colors exhibit higher saturation (s= 0.27-0.37) compared to transparent isotropic assemblies.

**Figure 5.**
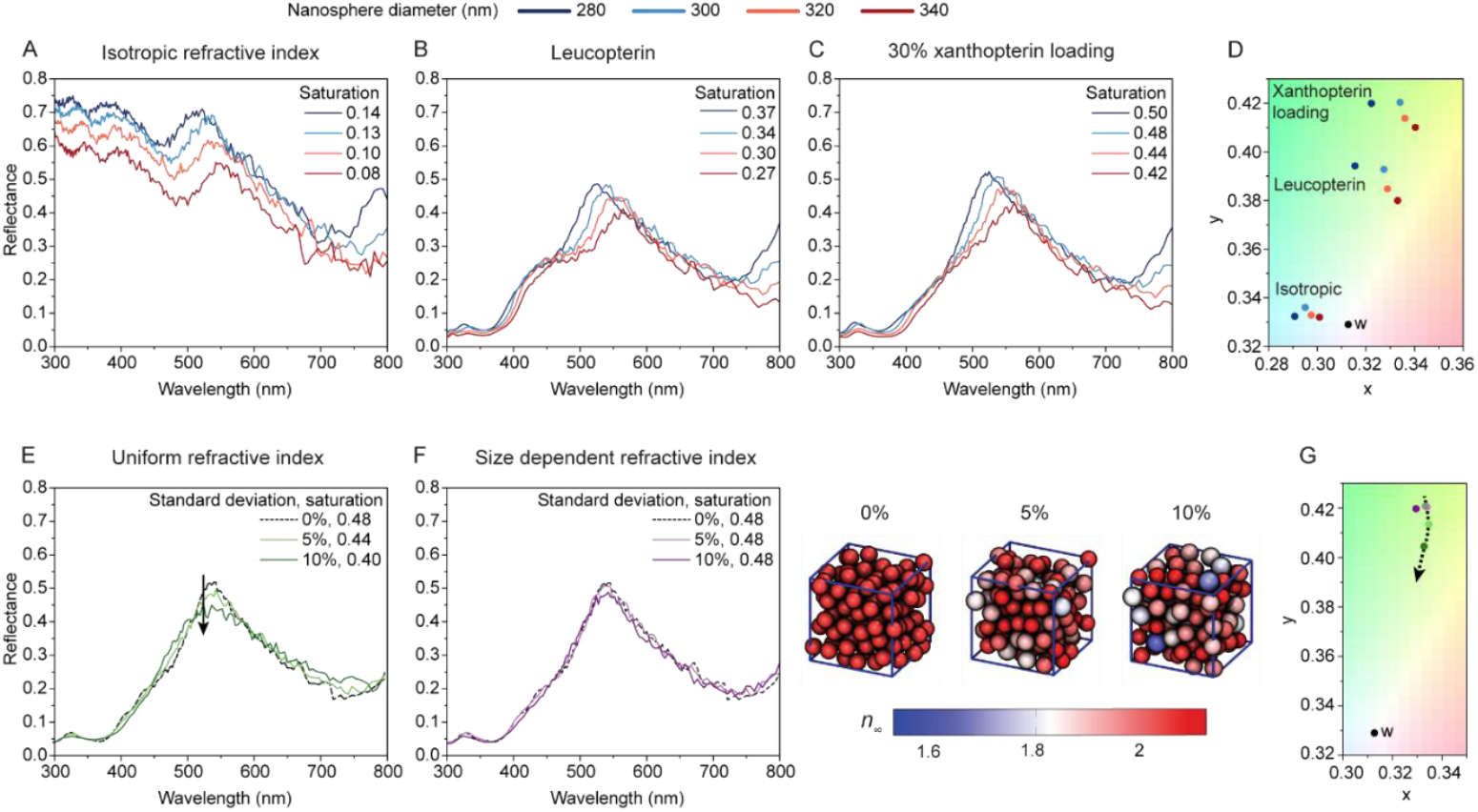
**(A-C)** FDTD simulated reflectance spectra of MD generated 5×5×5-*μ*m^3^ photonic glass slabs comprising nanospheres with 5% standard deviation, 55% filling fraction, varying diameters, size dependent refractive index and (A) isotropic or (B-C) wavelength-dependent complex refractive index (B; leucopterin nanospheres, C; 30% xanthopterin, 70% leucopterin nanospheres). **(D)** The reflectance spectra in (A-C) plotted on a CIE chromaticity diagram exhibiting the increasing saturation of the reflected colors. **(E-F)** FDTD simulated reflectance spectra of MD generated 5×5×5-*μ*m^3^ photonic glass slabs comprising nanospheres of 300 nm average diameter, 55% filling fraction, varying % standard deviation and (E) a uniform refractive index or (F) size dependent refractive index. (F, Right); snapshots of 1.5x1.5x1.5-*μ*m^3^ subregions of the MD-simulated particle configurations in (F) with color indicating the *n*∞ value used to calculate refractive index. **(G)** The reflectance spectra in (E-F) plotted on a CIE chromaticity diagram. Black arrows in (E) and (G) indicate the direction of increasing polydispersity and decreasing saturation. Saturation values of all spectra are specified in the legends. w; white point of standard illuminant D65 corresponding to average daylight.

To investigate the role of xanthopterin, a yellow pigment with strong visible absorbance (λ_max_ =388 nm) and significant refractive index dispersion (SI Appendix, Fig. S5A, S6B), we simulated assemblies composed of 30% xanthopterin and 70% leucopterin (vol. fraction) (Fig. 5C). The complex refractive index of pure xanthopterin was calculated as described for leucopterin (SI Appendix, Fig. S6B), and that of the mixture was calculated using the Lorenz-Lorenz mixing rule (eq. 3-8, SI Appendix, Fig. S7) (59). The reflectance spectra exhibit the same peak positions as the leucopterin assemblies, but with a significantly sharper reflectance peak and thus, a marked increase in color saturation (Fig. 5D) (s= 0.42-0.50). This phenomenon arises from the interdependent effects of pigment absorption (dampening of the shoulder peak at ∼445nm) and anomalous refractive index dispersion which enhances structural peaks in the green (33).

To investigate how the refractive index-size dependency (Fig. 3D) influences coloration, we compared simulations of 300 nm (average diameter), xanthopterin-loaded assemblies with varying polydispersity (0-10%) where the particles (i) all have the same complex refractive index (Fig. 5E) and (ii) where refractive index depends on size (Fig. 5F). For (i), the reflectance peak broadens as the polydispersity increases - an expected outcome of increased disorder, reducing saturation from 0.48 to 0.40 (Fig. 5E). In contrast, no reflectance broadening is seen when particles have a size-dependent refractive index, and the saturation remains high (s= 0.48) even with a large polydispersity of 10% (Fig. 5F). The same trend was observed for all nanosphere sizes between 280-340 nm (SI Appendix, Fig. S8). This effect likely occurs because the inverse linear relationship between nanosphere refractive index and diameter causes the optical path length (OPL = n*d; n=refractive index, d= physical distance light travels in the medium) of the nanospheres to become more uniform and thus reinforces the short-range order (structure factor (21, 60)) in the assemblies. Size dependent refractive index thus mitigates against the dampening effect of polydispersity and allows the system tolerate distributions of particle sizes while maintaining highly saturated colors.

## Discussion

The application of photonic glasses as alternatives to conventional absorbing dyes and iridescent materials is limited by their low color saturation (20–23). Here we show that the vivid colors of damselflies are tuned *via* precise modulation of the size of pteridine nanospheres in a photonic glass. Color saturation is enhanced by two ‘ingenious’ strategies: First, doping of transparent nanospheres with xanthopterin, a yellow pigment, dramatically enhances saturation by the absorption of blue-wavelengths coupled with refractive index (material) dispersion near the absorption band due to Kramers-Kronig relations (33, 61). Sai *et al*. (28) predicted theoretically that loading polystyrene particles with beta carotene can expand the color gamut and saturation of disordered photonic assemblies. However, to the best of our knowledge, this is the only known ‘real’ system employing narrowband pigment loading *within* colloidal particles. The scarcity of synthetic examples utilizing this powerful strategy is likely due to the challenges of synthesizing heterogeneous composite particles. Second, the dependency of refractive index on particle size (structural dispersion), which occurs through tuning of the crystallinity of the particles as a function of size, introduces the system with a tolerance against reflectance broadening – maintaining high color saturation even in disordered, polydisperse systems.

Remarkably, the tuning of particle size, pigment loading and particle size-refractive index dependence arise spontaneously from nanosphere formation during ontogeny. As the thorax transforms from green to blue, densification of the nanospheres causes them to shrink, and their increasing crystallinity results in an increase in refractive index. The increase in absolute pteridine concentration, caused by ongoing pteridine biosynthesis in the nanospheres, appears to drive densification and crystallization. Pigment loading is also an inevitable consequence of pteridine biosynthesis - xanthopterin is a biosynthetic precursor to leucopterin (54–56) and will thus be naturally present in the nanospheres during leucopterin synthesis. The relative shift from xanthopterin to leucopterin in the thoraxes during development is also indicative of an ongoing maturing metabolic activity of the organism (45), which ultimately drives the color change, and saturation enhancement.

Interestingly, Zhang *et al*. (62) reported a similar process in lizards, where ontogenetic color change arises by a gradual maturation of an existing pigment cell rather than its replacement with new pigment cell type. Our observation in a phylogenetically distinct organism suggests this may be a widespread and general phenomenon underlying ontogenetic color changes in animals. While this study is on male blue-tailed damselflies, the same optical mechanism is likely responsible for violet, blue and green coloration in female morphs of blue-tailed damselflies and other species of the Odonata order (40–42). More broadly, biological photonic glass structures, whether pigment-loaded or purely dielectric, are also probably more widespread than currently known. They may occur in other insect orders, where pteridines are abundant and function as pigments and excretory substances (54), but likely extend across a wide range of taxa, morphologies and material compositions.

While the formation of guanine, the most abundant biogenic organic crystal, has been described in detail (63–69), this study presents the first insights into the formation of optically functional biogenic pteridine crystals (34, 48, 49). Our results indicate that the gradual onset of molecular ordering in the nanospheres occurs due to the increased pteridine biosynthesis and metabolic activity in the organelles, consistent with a non-classical crystallization mechanism (70). The absence of fully periodic crystals in the damselflies is likely due to the large number of different pteridines present in the organelles. Understanding how biology forms birefringent spherulites may inspire new strategies for improving the development of synthetic mimics to these materials - as alternatives to inorganic particles in photonic assemblies (71).

### Concluding remarks

We report two biological strategies for enhancing color saturation in photonic glasses that have not been conceived or practically manifested in synthetic optical devices. The degree of tolerance of color saturation towards variations in particle size is a strong demonstration of how biology’s complex and sophisticated solutions in optics offer new understanding in scattering physics that can inspire the development of efficient and sustainable organic replacements to conventional synthetic pigments. Several questions remain about how photonic structures are biologically controlled in organisms, including the degree of genetic regulation and the mechanisms of precursor synthesis or transport to the transforming tissues.

## Materials and Methods

### Specimen collection

Damselflies were collected during June-August 2024 in two locations in Israel (River Park, Beersheva and Ekron River, Mazkeret Batya). Use of insects for research purposes does not require animal ethics approval in Israel, but animals were handled with established best practices.

### Optical microscopy

A Zeiss Discovery.V20 stereomicroscope equipped with an Axiocam 305 color camera were used to take images of the damselflies.

### Microspectrophotometry

The reflectance was measured from the thorax area of each male damselfly on the day of collection or the following day while the damselflies were alive. To correlate between structural/chemical properties and damselfly color, each damselfly was labelled and stored for further analyses (electron microscopy or LCMS). Reflectance spectra were obtained using a Zeiss AX10 microscope equipped with a Zeiss X10/0.25 NA HD DIC objective. The specimens were epi-illuminated with an LEJ HXP 120-V compact light source. The reflected light was collected from the same objective into an optical fibre (QR450-7-XSR, Ocean Insight) and the spectra measured using an Ocean Insight FLAME miniature spectrometer. The spectra were normalized using a white diffuse reflectance standard (Labsphere USRS-99-010, AS-01158-060). Background spectra were measured when no light was applied. Reflectance spectra were obtained over the range of 180– 880 nm. The integration time ranged from 10 to 40 s, with 1-10 averaged scans for each measurement.

### CIE chromaticity diagrams

International Commission on Illumination (CIE) diagrams were created using Origin(Pro) (version 2023b, OriginLab Corporation, Northampton, MA, USA). The CIE 1931 color space and D65 illuminant (represents average daylight illumination) were used.

### Cryogenic-SEM sample preparation and imaging

On the day of collection or the following day, the area of interest (first section of the abdomen or tail) was dissected and chemically fixed with 4% paraformaldehyde (PFA, CAS 50-00-0, Thermo Scientific Chemicals) and 2% glutaraldehyde (GA, CAS 111-30-8, Sigma-Aldrich) in 1xPBS for 3-4 hours. The chemically fixed tissues were embedded in 7% agarose gel and sectioned to 180-micron thick slices using (LEICA VT1000 S Vibratome). The sections were sandwiched between two aluminum discs in a 15% dextran (CAS 9004-54-0, Sigma-Aldrich) in 1X PBS solution and cryo-immobilized in a high-pressure freezing device (EM ICE, Leica). The frozen samples were then mounted on a holder under liquid nitrogen in a specialized loading station (EM VCM, Leica) and transferred under cryogenic conditions (EM VCT500, Leica) to a sample preparation freeze-fracture device (EM ACE900, Leica). Samples are fractured at −120 °C by hitting the top disc carrier with a tungsten knife at a speed of 135-140 mm/s, this exposed a clean fracture plane which can be imaged. The samples were etched at −110 °C for 3:00-3:40 min to evaporate excess water, and coated with 3 nm of Pt/C. The samples were imaged in an HRSEM Gemini 300 scanning electron microscope (Zeiss) by a secondary electron in-lens detector while maintaining an operating temperature of −120 °C.

Filling fraction (ff) was estimated from cryo-SEM images by measuring the 2D area of nanospheres in cells and dividing by the total area of interest. Assuming the sample is isotropic, and particles are roughly spherical, then 2D ff ≈ 3D ff.

### Nanosphere extraction for TEM imaging

The nanospheres were extracted from the thorax area by washing the specimen with ultrapure water and dissecting the colored areas into a vial with hexane or ethanol. The vials were sonicated briefly to release the nanospheres. The resulting suspension was drop-casted on a carbon-coated Cu-meshed TEM grid and allowed to dry.

### TEM imaging (measuring nanosphere diameters) and electron diffraction

Extracted nanospheres were imaged with a ThermoFisher Scientific (FEI) Tecnai T12 G2 TWIN TEM operating at 120 kV. Images and electron-diffraction patterns were recorded using a Gatan 794 MultiScan CCD camera. Electron-diffraction analysis was done using Gatan DigitalMicrograph software with the Diffpack module. Nanosphere diameters of samples with different colored thoraxes (reflectance from the thorax was measured as described above) were measured from TEM images of extracted nanospheres using Gatan DigitalMicrograph software. Each sample was extracted from an individual damselfly, 70-100 nanospheres were measured.

### Refractive index measurements by front scattering experiments

Scattering measurements of single nanospheres were performed using a commercial inverted optical microscope (Zeiss, Axio Observer 5) conjugated with a custom-built optical setup. The sample was illuminated with a ‘white’ light-emitting diode in a dark-field configuration and collected using a low NA objective to reduce unscattered light collection (N-Achroplan, ×20, NA.0.45). The light was then directed outside the microscope into the additional setup that further magnified the image (×6) and coupled into a fiber (105–150 μm in diameter, depending on nanospheres size) connected to a spectrometer (Shamrock 303i, Andor). The nanospheres were illuminated using an annulus (Θi = 25°). Remaining unscattered light was filtered out using an iris placed at the Fourier plane (imaged by a Blackfly S USB3, FLIR). The experimentally obtained spectrum was normalized according to the background signal and source spectrum and was then compared with refractive index dependent calculations of the scattering cross-section to extract the refractive index. The calculations were done using MATLAB according to the dimension or each nanosphere as extracted from TEM images of the specific measured nanosphere using ImageJ. Blurred edges of certain nanospheres due to remains of organic residues around them may have resulted in an underestimation of their diameters, leading to an overestimation of their refractive index as discussed in the main text. Detailed information about the optical setup and calculations of the scattering cross-section is available in ref. (51).

### Pteridine pigments extractions

Damselfly specimens were euthanized by plunge freezing in liquid nitrogen on the day of collection or the following day after measuring their reflectance. The specimens were stored at -80°C until the extraction. All solvents used for extractions were of LC-MS grade. The sample set included 45 damselflies (45 thoraxes and 35 tails, some tails were used for other analysis or lost) which were randomized and extracted in batches on six separate days. The extraction method was developed for this experiment, partially based on refs. (72, 73). The thorax and tail were dissected and weighed separately using a Radwag XA 6.4Y.M PLUS Microbalance. Each specimen was transferred to a labelled 2 mL Eppendorf vial and 0.5 mL 1%NH_4_OH was added. Small scissors were used to lyse the tissue. Each sample was spiked with 5 μL of the 100 μg/mL internal standard biopterin-d3 (CAS 1217838-71-5, Santa Cruz Biotechnology). The samples were sonicated for 5 minutes. 0.5 mL methanol was added. The samples were incubated at -20°C for 60 minutes. Next, the samples were sonicated for 2 minutes and filtered through a 0.45-micron syringe nylon filter (424-FNY422013, Jet Biofil) into clean 2 mL Eppendorf vials. 100 μL 1% formic acid and 0.9 mL of methanol were added. The samples were fully dried in a SpeedVac and the pellets were stored at -80°C until the LC-MS analysis. The samples were resuspended on the day of LC-MS analysis in 50 μL 1% NH_4_OH and 60 μL acetonitrile (ACN), sonicated for 5 minutes and vortexed briefly. Then 10 μL of 1% formic acid was added to neutralize the pH. The samples were centrifuged and decanted twice, then transferred into LC vials for analysis.

### Liquid chromatography-mass spectrometry (LC-MS)

The commercially available analytical standards of the following pteridines were used to build a calibration curve for the analysis: isoxanthopterin (CAS 529-69-1, Santa Cruz Biotechnology), xanthopterin (CAS 5979-01-1, Santa Cruz Biotechnology), pterin (CAS 2236-60-4, Sigma-Aldrich), leucopterin (CAS 492-11-5, Biosynth), 7,8-dihydroxanthopterin (CAS 1131-35-7, BOC Sciences), biopterin (CAS 22150-76-1, AA Blocks), 7,8-dihydrobiopterin (CAS 6779-87-9, Aaron Chemical), pterin-6-carboxylic acid (CAS 948-60-7, Sigma-Aldrich) and sepiapterin (CAS 17094-01-8, A2B Chem). Biopterin-d3 (CAS 1217838-71-5, Santa Cruz Biotechnology) was used as internal standard.

To identify which pteridines are present in damselflies, we extracted pteridines from a set of damselflies with varying thorax colors and searched for masses of pteridines involved in animal coloration that were selected based on previous reports (SI Appendix, Table S1, refs. (1-11)). The masses detected in the samples matched 12 pteridines from the list. We then used the available commercial standards to confirm the identification of 7 of the pteridines in our test samples (marked with an asterisk in SI Appendix, Table S1). Since the solubilities of most pteridines are very low and some are almost insoluble, the concentrations of the stock solutions and of the calibration samples varied for the different pteridines. All solvents used were of LC-MS grade. We first prepared a concentrated stock solution of each compound by weighing the compounds using Radwag XA 6.4Y.M PLUS Microbalance (±0.007 mg) and dissolving them in 1% or 2% NH_4_OH. The solutions were sonicated until dissolved. The stock solutions were prepared by diluting the concentrated stocks 100X with ultrapure water and ACN to a final ratio of 1:1. We then prepared a master stock which was diluted to create six calibration samples. The master stock was created by adding all the standards to one vial and adding water and ACN to a final ratio of 1:1 as specified in SI Appendix, Table S2. The calibration samples were created by diluting the master stock, adding 5 μL of internal standard (biopterin-d3) and adding water and ACN to a final ratio of 1:1. The final concentrations of the calibration curve are specified in SI Appendix, Table S3. The calibration samples were stored at -80°C.

Pteridines were analyzed using Waters ACQUITY UPLC I-Class Plus System coupled with Thermo Exploris 240 high-resolution mass-spectrometer. A separation of the selected pteridines was achieved on Waters BEH Amide column (1.7 µm, 2.1 x 100 mm) using the following gradient of the mobile phase A (10mM ammonium formate in 10% acetonitrile, pH 4.0) and mobile phase B (10mM ammonium formate, pH 4.0 in 90% acetonitrile): 0-2 min 5% A, 2-15 min 50% A, 15-15.5 min 80% A, 15.5-17 min 80% A, 17-17.5 min 5% A, 17.5-20 min 5% A. The injection volume was 5 µL, the column temperature was kept at 30 °C, and the flow rate was 0.25 mL/min. The total run time was 20 min.

High-resolution mass spectra were acquired using electron spray ionization (ESI) in positive full scan mode (70-800 m/z) with a resolution 24000 full width at half-maximum (FWHM). The MS parameters (ion spray voltage, sheath gas, aux gas, sweep gas, ion transfer tube temperature, and vaporizer temperature) were set up according to the manufacture recommendations. Data was acquired under the control of Xcalibur software, v3.5 (Thermo Scientific). Xcalibur software was also used for the data analysis to extract targeted ions (EIC) (74).

The concentration of each compound was calculated using the calibration curve built using Quan browser for Xcalibur. The obtained results were normalized (divided by) to the sample weight, Log10transformed and subjected to K-means cluster analysis. The optimal number of clusters (two) was found using the Partitioning Around Medoids algorithm using R software (75) (SI Appendix, Fig. S9). K-means cluster analysis (76) was performed using Origin(Pro) (version 2023b, OriginLab Corporation, Northampton, MA, USA.). The initial cluster centers were not specified, and the maximum number of iterations was 10.

### Optical constants calculation using Kramers–Kronig Relation

The real (*n*) and imaginary (κ, extinction coefficient) parts of the complex refractive index (*n*+iκ) of leucopterin and xanthopterin were calculated from their absorbance spectra using the Kramers-Kronig relations. Absorbance was measured using a UV-visible spectrophotometer (Thermo Fisher Scientific EVvolution 220). Leucopterin (CAS 492-11-5, Biosynth) and xanthopterin (CAS 5979-01-1, Santa Cruz Biotechnology) powders were dissolved in 1% NH_4_OH and filtered with a 0.22 μm mesh filter. The absorbance was measured between 190-1000 nm. All calculations were performed using a custom Python script (Python 3.11, NumPy, SciPy, Matplotlib, and Pandas libraries). The code was developed with the aid of Gemini (Google) and ChatGPT (OpenAI).

To estimate the imaginary part of the refractive index (κ), the absorbance spectrum was normalized such that the maximum absorption coefficient (α) in the range of 320–700 nm matched a target value of α_max_=2.5×10^6^ m^−1^ (2.5 μm^−1^). α_max_ was estimated based on measurements from ref. (61). The absorption coefficient α(λ) was then converted to κ(λ) using the relation:

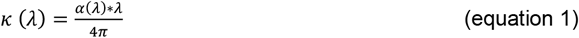

Where λ is the wavelength in meters.

The real part of the refractive index (*n*) was computed via the Kramers–Kronig relation (57):

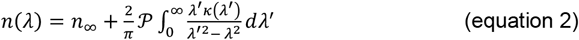

Where *n*∞ is the refractive index at infinite wavelength (a constant representing the electronic contributions at very high frequencies, or essentially the refractive index far from any absorption bands), 𝒫 denotes the Cauchy principal value of the integral. The integral was approximated numerically using the trapezoidal rule. *n*∞ values were estimated based on the experimental measurements of the tangential refractive index of the nanospheres, *n*_t_. To account for the refractive index varying with nanosphere diameter, a linear fit of the experimental measurements of *n*_t_ (SI Appendix, Fig. S4A) was used to estimate a range of *n*∞ values corresponding to increasing nanosphere diameters (SI Appendix, Fig. S4B). The real part of the refractive index, *n*, was computed for each *n*∞ value.

### Calculation of the complex refractive index for a two-component mixture

To compute the complex refractive index spectra of a two-component mixture, we applied the Lorentz–Lorenz mixing rule (59) to the complex dielectric permittivity of each component. The real (*n*) and imaginary (κ) components of the refractive index were obtained for each pure compound over 190-1000 nm wavelength range as previously described. The same value of *n*∞ was used for the two components in the mixture. The real (*n*) and imaginary (κ) components of the refractive index were computed for each *n*∞ value corresponding to varying nanosphere diameters.

The complex refractive index N=*n*+iκ was squared to obtain the complex dielectric function ϵ=N_2_. For each pure component (denoted as 1 and 2), the Lorentz–Lorenz transformation of the dielectric function was computed as:

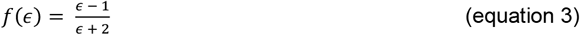

The effective Lorentz–Lorenz factor of the mixture was calculated via a linear volume-fraction-weighted average:

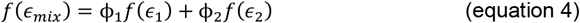

where ϕ_1_ and ϕ_2_=1−ϕ_1_ are the volume fractions of the two components.

This expression was inverted to solve for the complex dielectric function of the mixture:

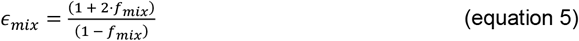

Finally, the complex refractive index of the mixture was obtained from the square root of the dielectric function:

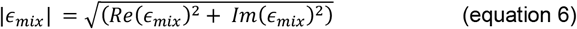

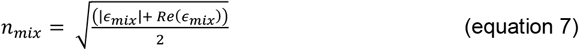

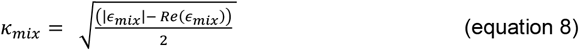

To ensure physical validity, the terms inside the square roots are clamped to be non-negative.

### Saturation of color

Saturation was calculated as in ref. (77):

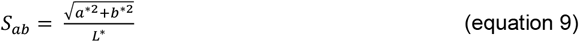

Where L* represents the lightness, and a*, b* are the chromaticity coordinated in CIELAB space. All reflectance spectra were normalized (divided by the maximum value) before the coordinates were calculated so the simulations and experimental data are comparable.

### Simulated reflectance spectra

MD simulations were carried out in similar manner to ref. (5) using HOOMD-blue (version 2.9.4) (78), with the Langevin integrator (*kT* = 0.05) and repulsive polydisperse 12–0 (LJ 12-0) pair-potential:

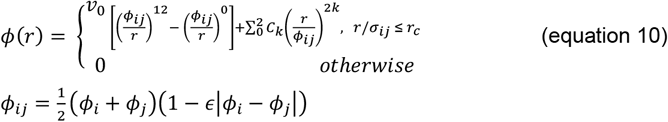

where *v*_0_ = 1, *ϵ*_*ij*_ = 0.01, *σ*_*a*_ is the diameter of particle *a*, and *r*_c_ = 1.5*μ*_1_ is the cutoff distance, using the PolydisperseMD(79) plugin. Particle sizes were randomized from a log-normal distribution, with mean 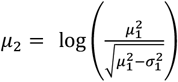 and 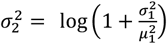, and the values outside range [100nm, 600nm] were rounded to the nearest boundary to avoid the chance of extreme particle sizes. In the simulations, mean particle sizes *μ*_1_ = [280nm, 300nm, 320nm, 340nm] and standard deviations of *σ*_1_= [0, 5%, 10%] were used. Simulations were initiated with random placement of particles in a 20 × 20 × 20-μm^3^ box with periodic boundary conditions (PBC). Particle overlaps were relaxed with repulsive gaussian potential for 2000 time steps, and the systems was then squeezed to a final 5 × 5 × 5-*μ*m^3^ volume during the rapid quench, in 5 × 10^5^ time steps while using the LJ12-0 pair potential. Particle coordinates and diameters were then imported to Lumerical FDTD (Ansys) release 2025 R1.3. In the FDTD simulation set-up, PBC boundaries in the *x* and *y* directions were used and a plane-wave source and reflection monitor were placed 6.5 and 7.5 *μ*m above the sample (in the *z* direction), respectively. The recorded reflectance was integrated in Lumerical over the 5 × 5-*μ*m^2^ monitor area.

## Data, Materials, and Software Availability

All data are available in the main text, in the Supplementary Information, and experimental datasets will be provided or deposited in an online repository after publication.

Source code will be provided, or code will be available in an online repository after publication.

## Acknowledgments

We thank Uri Ben Nun for collecting the damselflies for this study. Funding was provided by a European Research Council Starting Grant (grant no. 852948, “CRYSTALEYES”), a Human Frontier Science Program grant (grant no. RGP0037/2022), an Israel Science Foundation grant (grant no. 1565/22) awarded to B.A.P, a European Research Council Advanced Grant BoX-BOOM (No. 101096020) awarded to D.O and by a Research Council of Finland grant 347789 awarded to J.S.H. In addition, B.A.P. is the Nahum Guzik Presidential Recruit and the recipient of the 2019 Azrieli Faculty Fellowship. D.O. is the incumbent of the Harry Weinrebe Professorial Chair of Laser Physics. L.A. acknowledges the support of the Tom and Mary Beck Center for Advanced and Intelligent Materials. T.L. acknowledges the support from the Azrieli Foundation for the award of an Azrieli Graduate Fellowship 2024/25. We acknowledge the computational resources provided by the Aalto Science-IT project.

