## Supplemental Information for "Damselflies Overcome Color Saturation Barriers of Photonic Glasses *via* Structural Dispersion and Pigment Loading"

**Supporting Information**

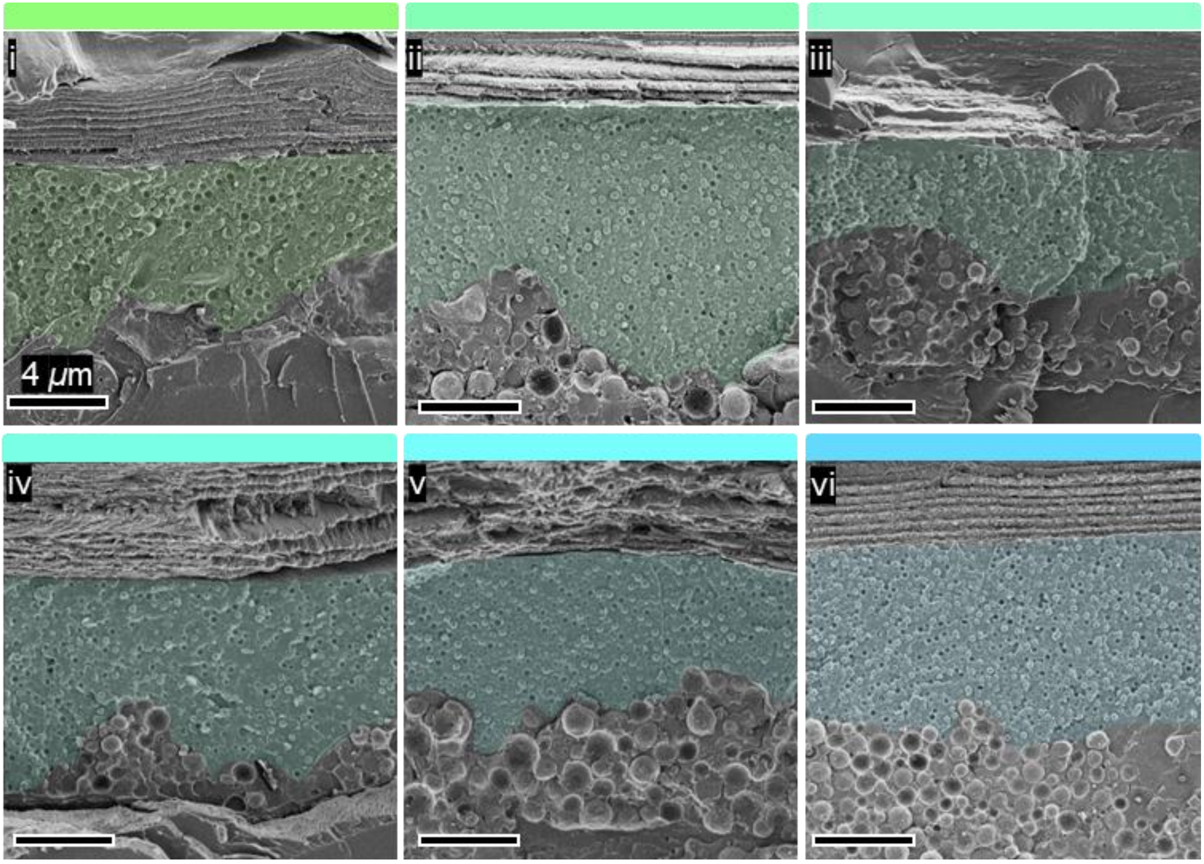


**Figure S1.** Cryo-SEM images of cross sections from thoraxes (i-v) and tail (vi) of the epidermis exhibiting the ultrastructure of differently colored damselflies. The distal epidermis containing the pteridine nanospheres is pseudo colored. Color panels; CIE representation of the reflectance measured from the specific tail/thorax which was analyzed.

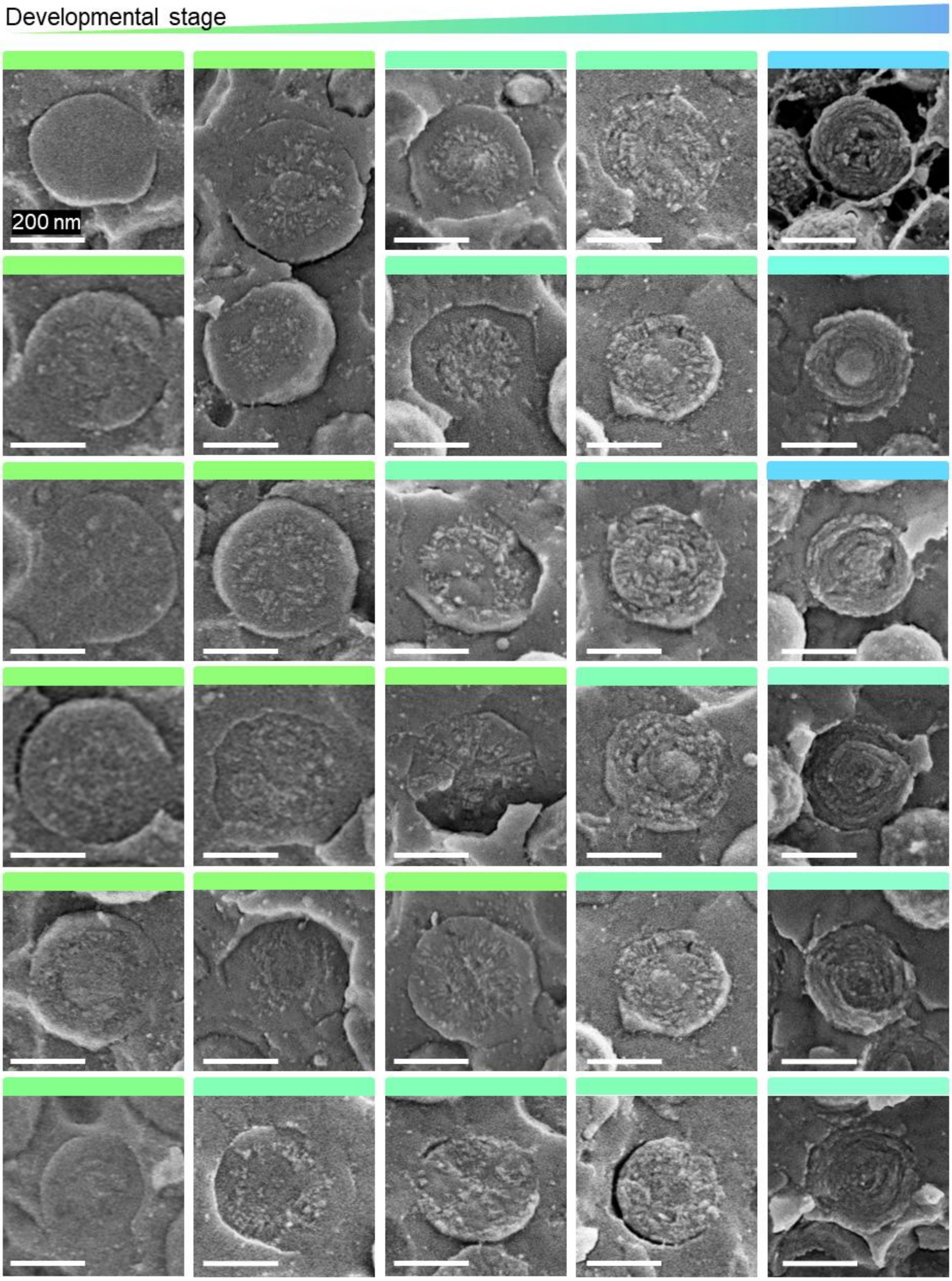


**Figure S2.** Cryo-SEM images of nanosphere cross-sections from damselfly samples of different colors. Color panels; CIE representation of the reflectance measured from the specific tail/thorax which was analyzed.

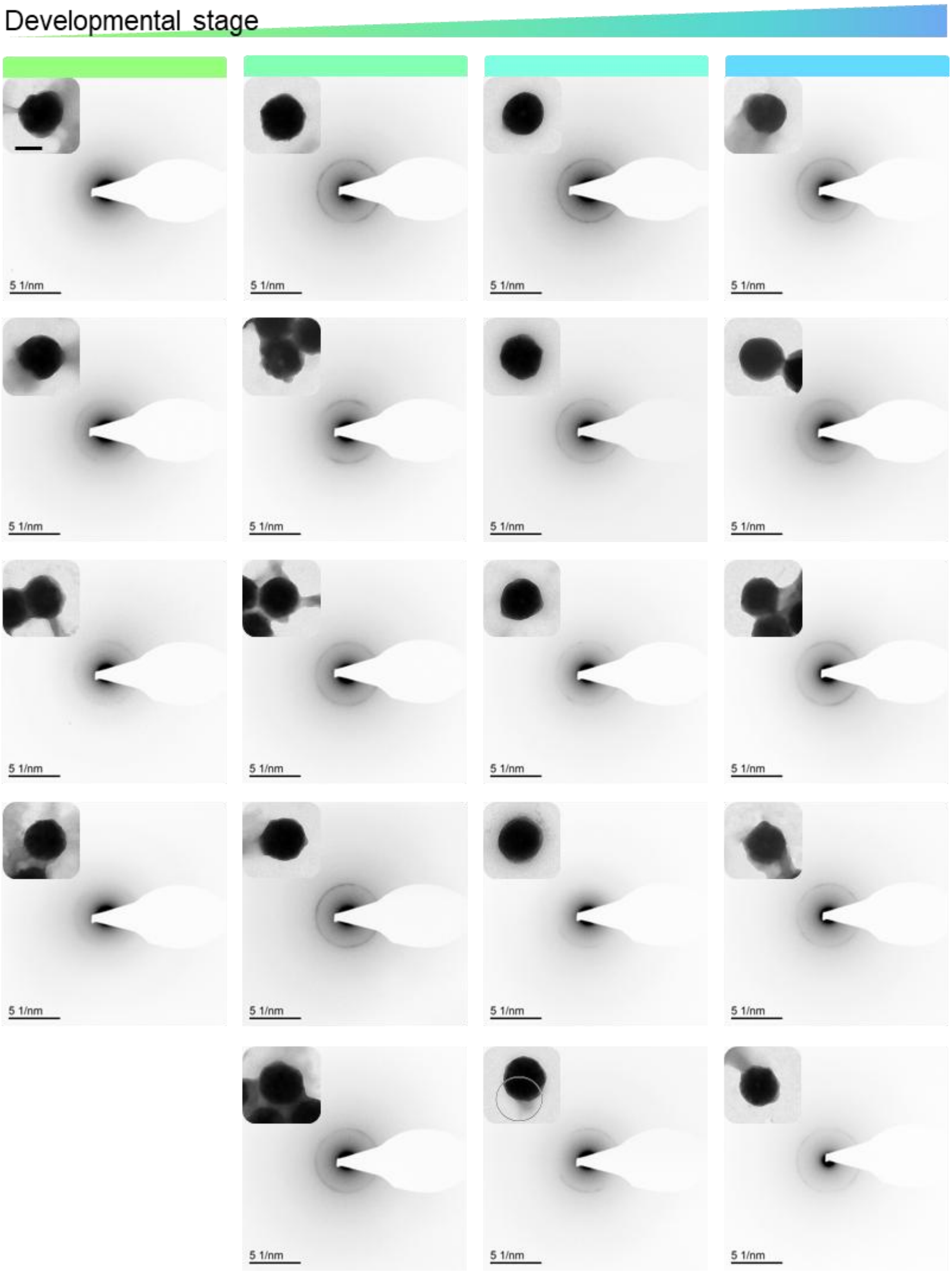


**Figure S3.** Electron diffraction patterns obtained from nanospheres extracted from damselfly samples of different colors. Color panels; CIE representation of the reflectance measured from the specific tail/thorax which was analyzed in each column.

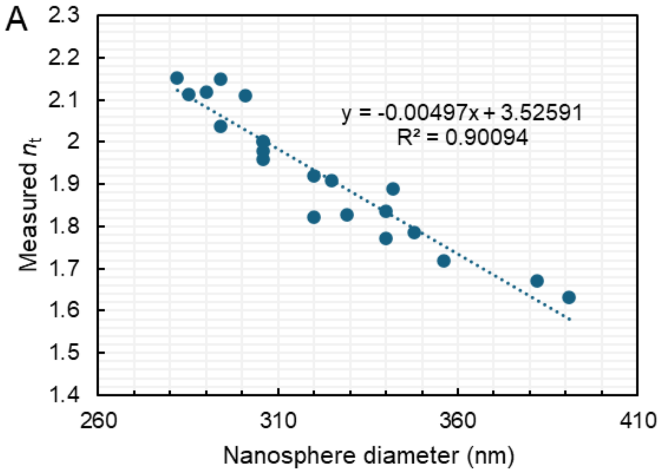


B

| Diameter (nm) | Corresponding $n_\infty$ |
|---|---|
| 280 | 2.126 |
| 290 | 2.076 |
| 300 | 2.026 |
| 310 | 1.976 |
| 320 | 1.926 |
| 330 | 1.876 |
| 340 | 1.826 |
| 350 | 1.776 |
| 360 | 1.726 |
| 370 | 1.676 |
| 380 | 1.626 |
| 390 | 1.576 |
| 400 | 1.526 |

**Figure S4. (A)** Linear fit of experimental measurements of $n_t$. Two outlying data points were not incorporated into the fit. **(B)** Range of nanosphere diameters and their corresponding *n* (Fig. 5A) or $n_\infty$ (Fig 5B-C, E-F) values from the linear fit. The $n_\infty$ values were used to compute the refractive index of nanospheres with varying diameters in Fig 5B-C, E-F.

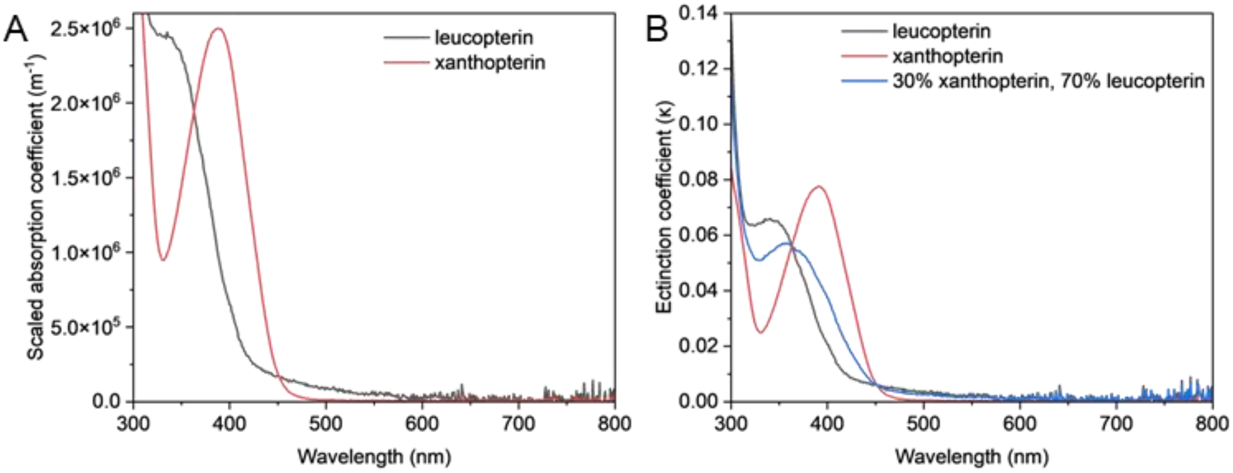


**Figure S5. (A)** Absorption coefficient of leucopterin and xanthopterin scaled to a maximum of 2.5e6 $m^{-1}$. **(B)** Calculated Imaginary component of the refractive index (extinction coefficient, κ).

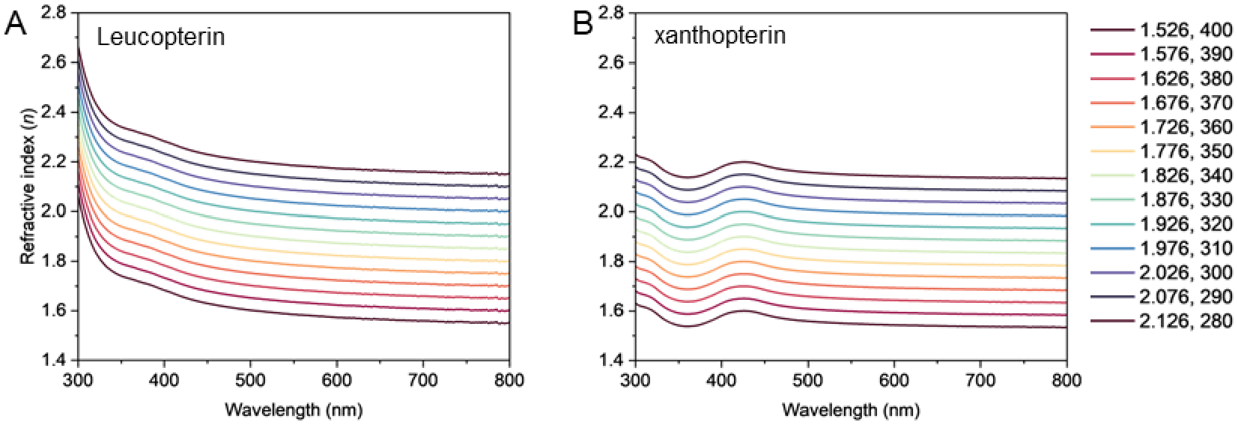


**Figure S6.** Calculated real component of the refractive index ($n$) with varying $n_\infty$ values of **(A)** leucopterin, **(B)** xanthopterin. Legend; $n_\infty$, corresponding nanosphere diameter for the simulations (nm).

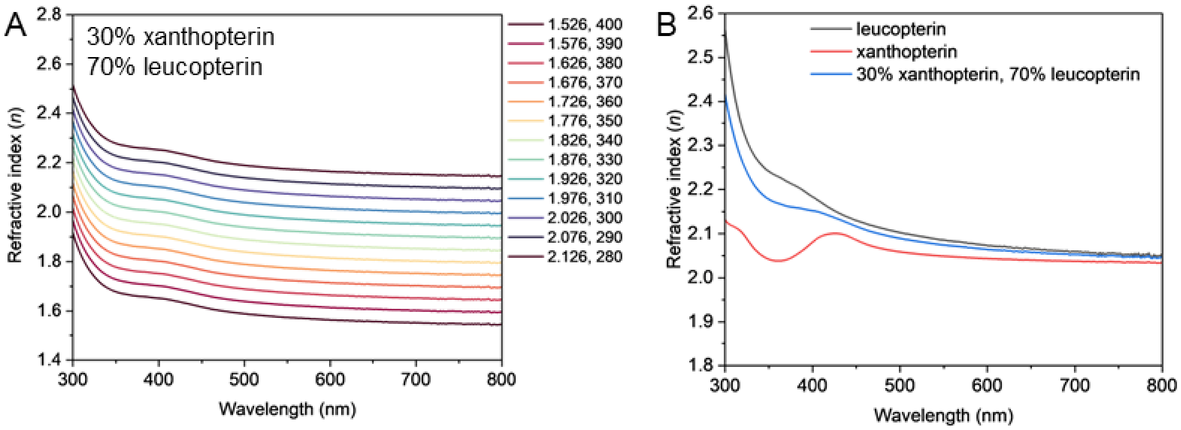


**Figure S7. (A)** Calculated real component of the refractive index ($n$) with varying $n_\infty$ values of a mixture of 30% xanthopterin, 70% leucopterin. Legend; $n_\infty$, corresponding nanosphere diameter for the simulations (nm). **(B)** Calculated real component of the refractive index with $n_\infty$=2.026 of leucopterin, xanthopterin and the mixture for comparison.

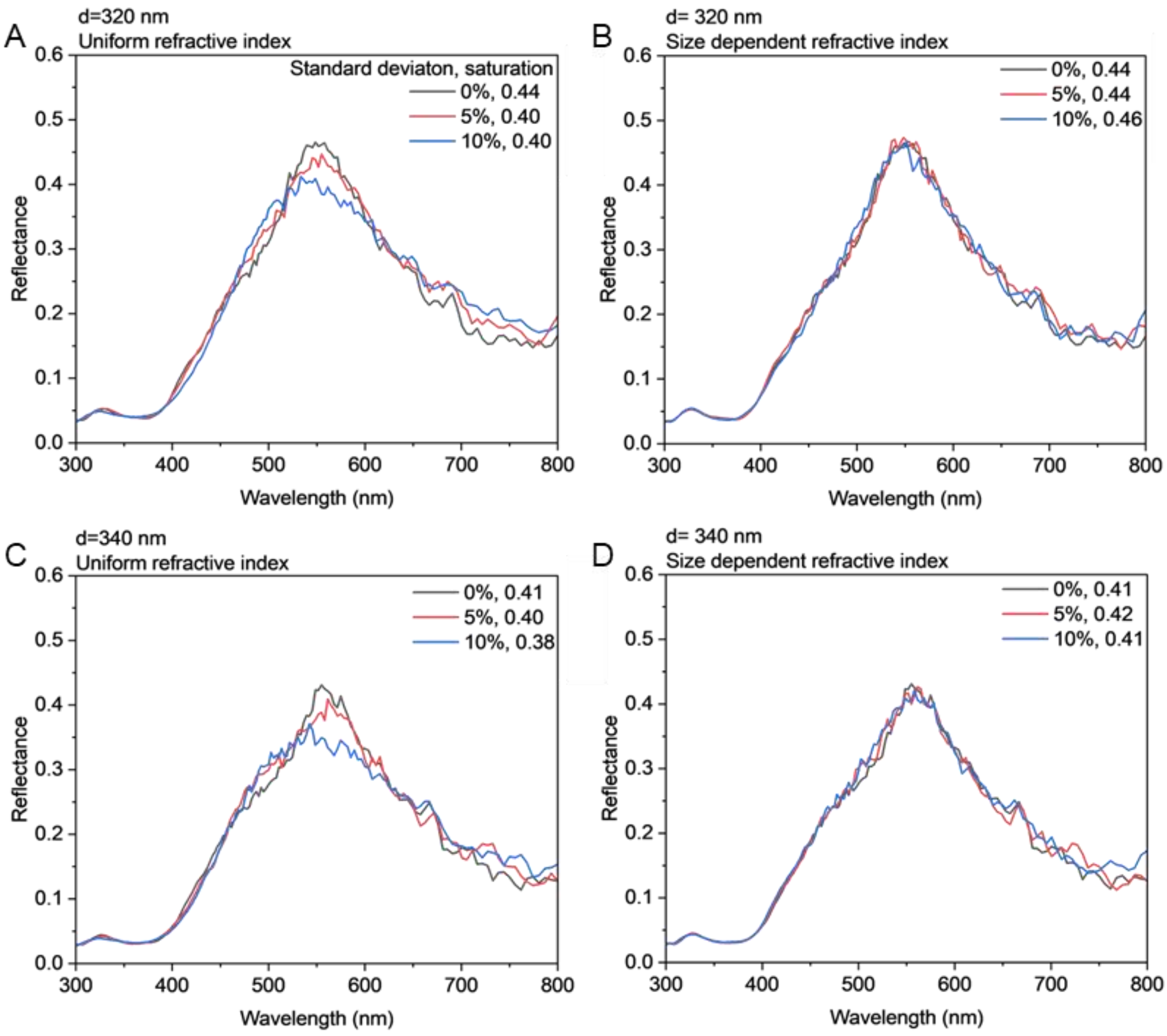


**Figure S8.** MD and FDTD simulated reflectance spectra of the 5×5×5-μm$^3$ photonic glass slabs comprising nanospheres of 320 and 340 nm average diameter, 55% filling fraction and varying % standard deviation.

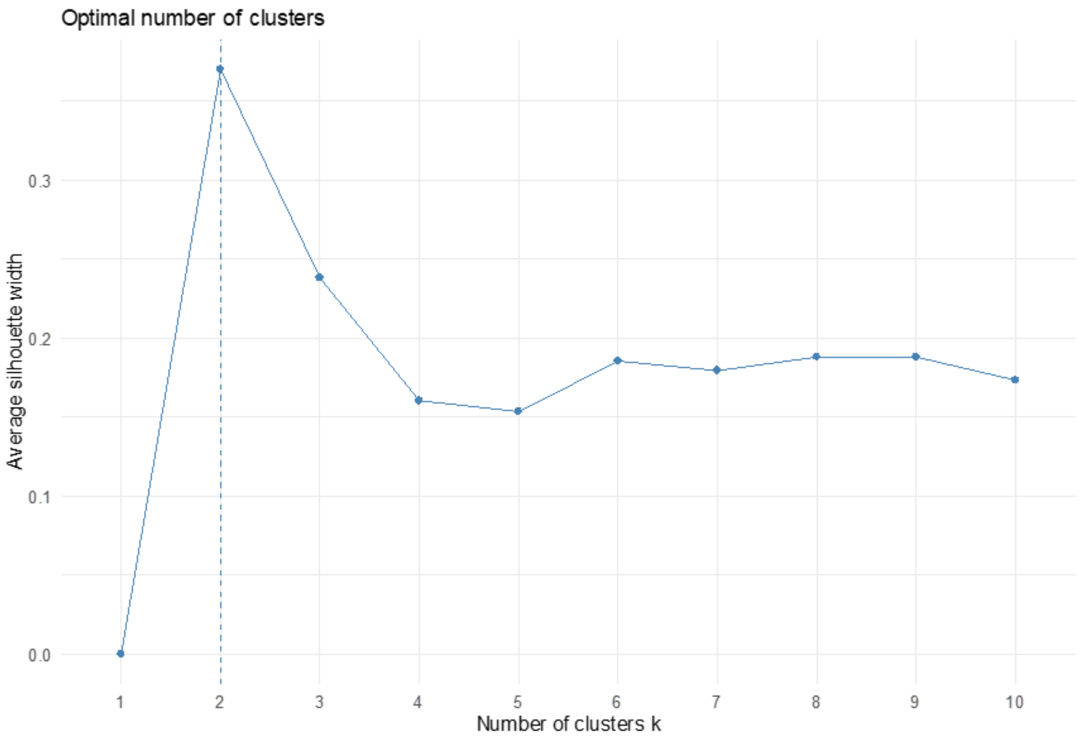


**Figure S9.** Elbow plot obtained from the Partitioning Around Medoids (PAM) method showing the optimal number of clusters in the LC-MS data is 2.

**Table S1. Pteridines involved in animal coloration based on previous reports**

| | **Formula** | **m/z** | **Retention time** |
|---|---|---|---|
| [*]Isoxanthopterin(1, 2) | C6H5N5O2 | 180.0516 | 3.9 |
| [*]Xanthopterin(1, 2) | C6H5N5O2 | 180.0516 | 3.9 |
| [*]Pterin(3) | C6H5N5O | 164.0567 | 2.71 |
| [*]Leucopterin(4) | C6H5N5O3 | 196.0465 | 5.62 |
| [†]Erythropterin(4) | C9H7N5O5 | 266.0519 | detected |
| [*]7,8-dihdroxanthopterin (5) | C6H7N5O2 | 182.0672 | 4.32 |
| [‡]Sepiapterin(1, 3) | C9H11N5O3 | 238.0935 | 2.73 |
| [*,‡]Biopterin(2, 3) | C9H11N5O3 | 238.0935 | 2.73 |
| [§]7,8-dihydrobiopterin(3) | C9H13N5O3+Na | 262.0911 | 5.45 |
| Neodrosopterin(1–3) | C15H16N10O4 | 401.1429 | x |
| Drosopterin(1, 2) | C15H16N10O2 | 369.1530 | x |
| Isodrosopterin(1, 2) | C15H16N10O2 | 369.1530 | x |
| Aurodrosopterin(2) | C15H15N9O3 | 354.1421 | x |

| | | | |
|---|---|---|---|
| 7-methylxanthopterin(6, 7) | C7H7N5O2 | 194.0673 | x |
| *Pterin-6-carboxylic acid(8) | C7H5N5O3 | 208.0465 | 1.73 |
| 2-amino-4-hydroxy-6-hydroxymethyl pteridine(1) | C7H11N5O8P2 | 356.0156 | x |
| | +Na | 377.9975 | x |
| Xanthurenic acid(2, 3) | C10H7NO4 | 206.0448 | x |
| Lumazine(9) | C6H4N4O2 | 165.0407 | x |
| 2-Amino-4-hydroxy-6-carboxypteridine(1) | C7H5N5O3 | 208.0465 | x |
| Deoxysepiapterin(3) | C9H11N5O2 | 222.0986 | x |
| ¶ Neopterin(7, 8) | C9H11N5O4 | 254.0884 | 3.53 |
| Isoxantholumazine (violapterin)(7, 10) | C6H4N4O3 | 181.0356 | x |
| Pterorhodin(4) | C13H10N10O4 | 371.0959 | x |
| Riboflavin(8) | C17H20N4O6 | 377.1456 | x |
| Isosepiapterin(11) | C9H11N5O2 | 222.0986 | x |
| 6-hydroxymethylpterin (ranachrome-3)(11) | C7H7N5O2 | 194.0673 | x |

* Compound was detected in damselflies, verified with a standard and quantified.

† No commercial standard, the mass was detected in very small abundance.

‡ Compound was not separated from its isomer (biopterin and sepiapterin)

§ Compound was detected in a very low concentration in damselflies, verified with a standard and quantified.

¶ Commercial standard was available but the compound was not detected in damselflies, the detected mass is probably an isomer of the compound.

**Table S2. Preparation of master stock**

| compound | Concentration of stock (ug/mL) | Volume of stock (mL) | concentration in master stock (ug/mL) |
|---|---|---|---|
| isoxanthopterin | 5 | 0.3 | 1.5 |
| xanthopterin | 25 | 0.06 | 1.5 |
| pterin | 12.5 | 0.12 | 1.5 |
| leucopterin (2% NH4OH) | 10 | 0.0140 | 0.14 |
| 7,8-dihydroxanthopterin | 1.6 | 0.08750 | 0.14 |
| biopterin | 100 | 0.02 | 2 |
| 7,8-dihydrobiopterin | 62.5 | 0.0304 | 1.9 |
| 3-Hydroxykynurenine | 62.5 | 0.0304 | 1.9 |
| Pterin-6-carboxylic acid | 0.1 | 0.3 | 0.03 |
| sepiapterin | 100 | 0.02 | 2 |
| | final volume | 1 | |
| | total volume of standards | 0.982300 | |
| | volume of ACN added | 0.015850 | |
| | volume of water added | 0.001850 | |

**Table S3. Concentrations of the calibration samples for LCMS analysis**

| | **Concentration (ug/mL)** | | | | | |
|---|---|---|---|---|---|---|
| **Compound** | 6 | 5 | 4 | 3 | 2 | 1 |
| isoxanthopterin | 1.5 | 0.75 | 0.15 | 0.075 | 0.015 | 0.0075 |
| xanthopterin | 1.5 | 0.75 | 0.15 | 0.075 | 0.015 | 0.0075 |
| pterin | 1.5 | 0.75 | 0.15 | 0.075 | 0.015 | 0.0075 |
| leucopterin | 0.14 | 0.07 | 0.014 | 0.007 | 0.0014 | 0.0007 |
| 7,8-dihydroxanthopterin | 0.14 | 0.07 | 0.014 | 0.007 | 0.0014 | 0.0007 |
| biopterin | 2 | 1 | 0.2 | 0.1 | 0.02 | 0.01 |
| 7,8-dihydrobiopterin | 1.9 | 0.95 | 0.19 | 0.095 | 0.019 | 0.0095 |
| 3-Hydroxykynurenine | 1.9 | 0.95 | 0.19 | 0.095 | 0.019 | 0.0095 |
| Pterin-6-carboxylic acid | 0.03 | 0.015 | 0.003 | 0.0015 | 0.0003 | 0.00015 |
| sepiapterin | 2 | 1 | 0.2 | 0.1 | 0.02 | 0.01 |
